# Genomic and transcriptional profiling stratifies VQ myeloma lines into two clusters with distinct risk signatures and drug responses

**DOI:** 10.1101/2022.08.21.504657

**Authors:** Evan Flietner, Mei Yu, Adhithi Rajagopalan, Yun Zhou, Yubin Feng, Anthony J. Veltri, Terra Lasho, Zhi Wen, Yuqian Sun, Mrinal M. Patnaik, Natalie S. Callander, Fotis Asimakopoulos, Demin Wang, Jing Zhang

**Author notes:** Corresponding authors: Demin Wang, Address: Blood Research Institute, 8727 Watertown Plank Rd Versiti, Milwaukee, WI 53226; Jing Zhang, Address: Wisconsin Institute for Medical Research II McArdle Lab for Cancer Research, 1111 Highland Avenue University of Wisconsin-Madison Madison, WI 53705. These authors contributed equally to this work (Evan Flietner, Mei Yu).

## Abstract

Multiple myeloma (MM) is a cancer of malignant plasma cells in the bone marrow and extramedullary sites. We previously characterized a VQ model for human high-risk MM. Different VQ lines display distinct disease phenotypes and survivals, suggesting significant intra-model variation. Here, we use whole exome sequencing and copy number variation (CNV) analysis coupled with RNA-Seq to stratify VQ lines into corresponding clusters: Cluster I VQ cells carried recurrent amplification of chromosome (chr) 3 and displayed upregulation of growth pathways and high-risk myeloma gene signatures, whereas Cluster II cells had monosomy chr5 and overexpressed genes and pathways associated with positive response to bortezomib (Btz) treatment in human MM patients. Consistently, in sharp contrast to Cluster II VQ cells that showed short-term response to Btz, Cluster I VQ cells were de novo resistant to Btz *in vivo*. Our study highlights Cluster I VQ lines as highly representative of human high-risk MM subset.

## Introduction

Multiple myeloma (MM) is a malignancy of terminally differentiated plasma cells (PCs) that primarily grow in the bone marrow (BM) [1]. MM arises from the pre-malignant condition monoclonal gammopathy of undetermined significance (MGUS), in which the accumulation of chromosomal copy number variations (CNVs), primary translocations, and somatic mutations leads to the expansion of a clonal PC population [2]. MGUS and MM share a number of overlapping CNVs, including hyperdiploidy [3], as well as primary translocations such as t(4;14) [4] and t(11;14) [5]. The genetic heterogeneity of MM is further increased by acquiring secondary CNVs [6] and point mutations [7] and a greatly altered landscape of DNA methylation compared to healthy PCs [8].

Genetic events (i.e. CNVs and primary translocations) present at time of diagnosis play a significant role in patient prognosis [9] and in the stratification of high-risk multiple myeloma (hrMM) [10, 11]. These events include two translocations involving the immunoglobulin heavy chain (IgH) locus: the t(4;14) translocation in which both fibroblast growth factor receptor 3 (FGFR3) and multiple myeloma SET domain (MMSET) are put under the control of the IgH promoter [12, 13], and t(14;16) in which the transcription factor c-MAF is overexpressed [14]. Amplification (≥4 copies) of the long arm of chromosome 1 (amp(1q)) is an hrMM prognostic marker [15], while a gain of a single copy (gain(1q)) is considered high risk when combined with a second hrMM chromosomal abnormality [16]. At the level of gene expression, two independent, and largely non-overlapping gene signatures predicting increased relapse risk and poorer overall survival have been developed for use in diagnosing hrMM: a 70 gene signature developed by the University of Arkansas for Medical Sciences (UAMS-70) [17], and a 92 gene signature developed by the Erasmus University Medical Center (EMC-92 or SKY92) [18]. Diagnosis of patients with hrMM is particularly important as these patients are likely to have a poor response to current treatment options [19, 20].

Murine models of MM play an essential role in dissecting mechanisms of disease growth [21] and as pre-clinical platforms for testing new anti-MM therapies [22]. The genetic basis for MM models ranges from the forced overexpression of a single MM-related oncogene such as the Eµ-MAF model [23], transgenic models in which oncogene overexpression is activated through plasma cell maturation as in the Vĸ*MYC model [24], or spontaneous MM development owing to genetic inbreeding as in the 5TMM family of myeloma models [25, 26]. Due to the heterogeneous genetic origins of human MM and the downstream pathological and therapeutic consequences, genomic characterization of these murine myeloma cells is necessary to establish what sub-type of MM they best model. Genetic landscape characterization of the 5TMM models has shown notable genomic differences between different lines, with the 5T2 line showing CNVs syntenic for gain(1q) MM whereas the 5T33 and 5TGM1 lines show CNVs syntenic for del(13q) [27]. Within transgenic MM models some chromosomal variability is also observed: CNV analysis of both primary and transplantable Vĸ*MYC showed that about 50% of sequenced lines had monosomy 5, and within these mice half additionally had monosomy 14, including a region syntenic to del(13) in human MM [28]. Further analysis of monosomy 14 Vĸ*MYC lines identified loss of the region containing the cell-cycle regulating miRNA cluster MIR15A/16-1 as driving MM progression in the Vĸ*MYC model and as a potential mechanism to explain del(13) as an early initiating CNV in the transition from MGUS to MM [29].

Previously, we developed and characterized a mouse model of MM driven by both the Vĸ*MYC transgene and oncogenic NRas^LSL-Q61R^ inducible via IgG1-Cre [30]. This so-called VQ model shows characteristics representative of hrMM, as it is highly proliferative, develops extramedullary disease, and is enriched for the UAMS-70 high-risk gene signature compared to control PCs [30]. In our initial study, five lines of VQ myeloma were isolated from primary mice (VQ-D1 through VQ-D5). In four of these VQ lines (VQ-D1, D2, D4, and D5), we established survival of recipient mice transplanted with donor VQ cells and carried out bulk RNA-Seq analysis. Finally, we established two VQ cell lines from two VQ-D2 recipients, termed VQ 4935 and 4938. Further drug testing of the VQ model also showed *de novo* resistance to the BCL-2 inhibitor venetoclax in the VQ cell lines, as well as reduced responses to the proteasome-inhibitors (PIs) bortezomib and carfilzomib in VQ-D1 recipient mice *in vivo* [31]. Although all VQ cells are driven by MYC dysregulation and RAS hyperactivation, different lines display distinct patterns of extramedullary disease, antibody sub-type, and survival (Figure 1A). The genetic and/or transcriptional bases for these phenotypic differences remain elusive. Additionally, what subtypes of hrMM they may individually represent (if any) are unknown. In the current study, we aim to address these questions through the molecular characterization of five VQ lines using B cell receptor (BCR) repertoire sequencing, whole exome sequencing, CNV analysis, and RNA- Seq.

**Figure 1.**
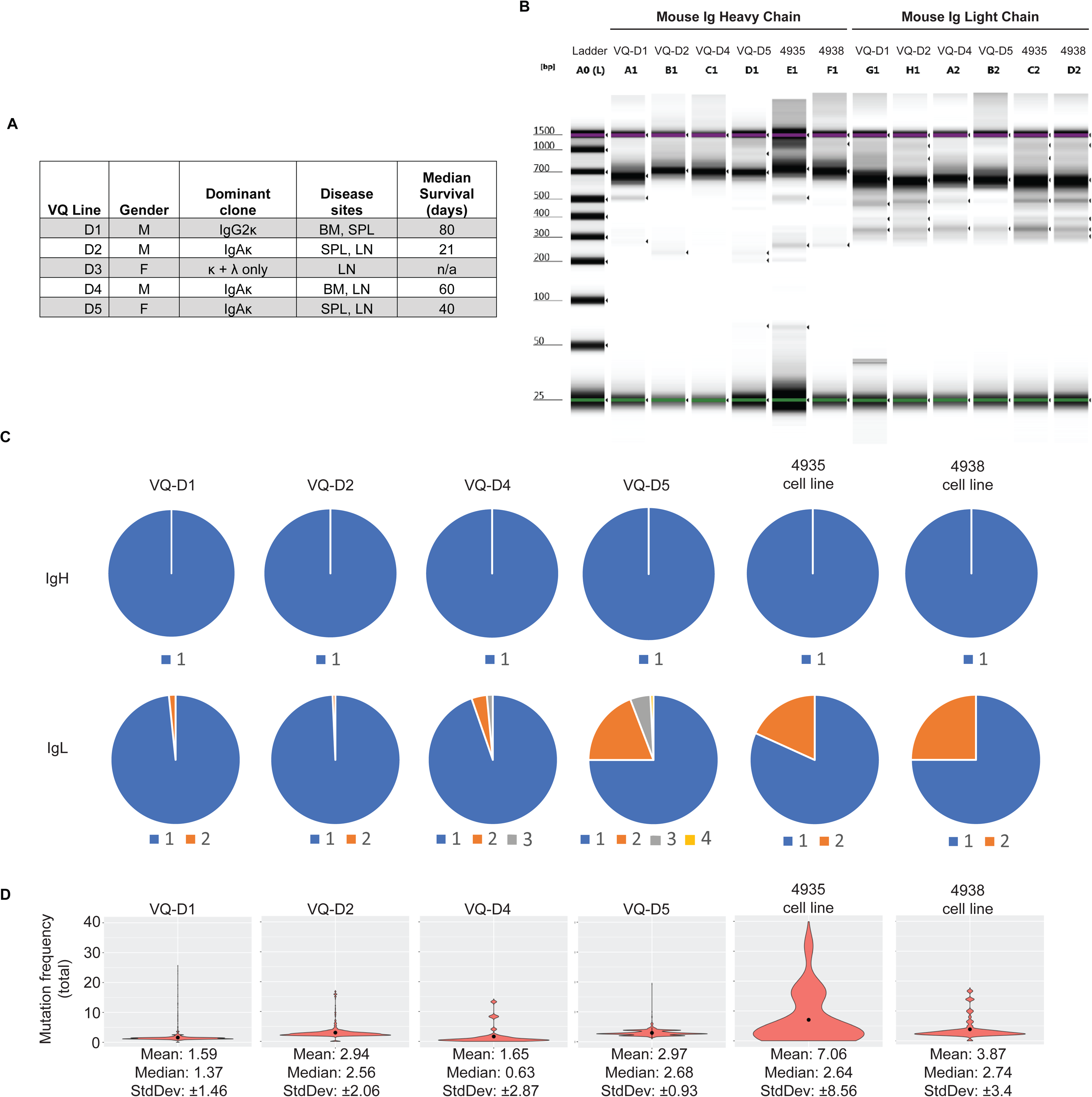
B cell receptor repertoire analysis shows dominant clonality and low somatic hypermutation (SHM) rates in primary VQ cells and VQ cell lines. (A) Table summarizing previously established characteristics of VQ donor lines. (B-D) B cell receptor heavy-chain and light-chain repertoire analysis was carried out as described in Methods. (B) Tapestation image showing immunoglobulin heavy chain and light chain library amplification for samples in panels C and D. (C) Pie charts depicting clonal frequency for immunoglobulin heavy chain (IgH) sequences (top) and immunoglobulin light chain sequences (bottom) sequences from primary VQ cells (VQ-D1, D2, D4, and D5) and from VQ-D2 derived cell lines (4935 and 4938). (D) Violin plots showing mean (black dots) and distribution of the total somatic hypermutation frequencies across B-cell IgH sequences from primary VQ cells (VQ-D1, D2, D4, and D5) and from VQ-D2 derived cell lines (4935 and 4938).

## Materials and Methods

### High-throughput sequencing of B cell receptor repertoire

We performed paired-end full-length high-throughput sequencing of the immunoglobulin heavey and light (IgH and IgL) chains using a unique molecular identifier (UMI)-based protocol detailed by Turchaninova et al [32]. In brief, we extracted mRNA, synthesized cDNA, and incorporated UMIs and a universal primer at the 5’ end with a template switch reaction. To prepare library, cDNA equivalent to 5 x 10^4^ B cells for each sample was amplified with two rounds of PCR. For immunoglobulin heavy chain amplification, the first PCR used a 5’ end universal primer and pooled 3’ end primers specific to mouse IgG or IgA constant region; for immunoglobulin light chain amplification, pooled 3’ end primers specific to mouse Igκ and Igλ were used. The second PCR used a 5’ and a 3’ universal primer with integrated sample barcoding at both ends. The Illumina adaptors were ligated to each sample library to allow for multiple sample sequencing simultaneously. We prepared barcoded libraries for each sample and then pooled the libraries to perform asymmetric 400 + 100 bp paired-end IgSeq in one run using Miseq Reagent Kit V3 (Illumina, Miseq Reagent Kit V3, 600-cycles) on Illumina MiSeq platform.

### B cell receptor repertoire analysis

Sequences were firstly demultiplexed and unique molecular identifier (UMI) were extracted by MIGEC v1.2.9 as previously described [32]. Briefly, we demultiplexed fastq files using sample barcodes and estimated the MIG (molecular identifier groups, a set of reads tagged with the same UMI) size distribution. Sequences with the same UMI in the presence of at least 2 times were then assembled and paired sequences were merged from 5’ end to 3’ end reads. We then mapped MIG consensus sequences to mouse IgH or IgL germline references and assembled clonotypes using MiXCR v3.0.3 [33]. Somatic hypermutation frequencies were calculated using the SHazaM package v1.1.0 [34] and plotted using ggplot2 v3.3.5 in R v4.0.3.

### Whole exome sequencing

Genomic DNA was extracted from CD138+ BM cells and tail tissues of moribund *V*ĸ*\*Myc; Nras^LSL-Q61R/+;^ IgG1-Cre* mice using Gentra Puregene Cell Kit (Qiagen). Whole-exome targeted capture was carried out using the SureSelect XT Mouse Exome Kit, 49.6 Mb (Cat# 5190-4641; Agilent Technologies). Exome capture, exome library amplification, and data analysis was carried out as described previously [35].

### Copy number variation analysis

Whole exome sequencing data was aligned to the mm10 reference genome using BWA MEM [36]. Copy number variation analysis was carried out using matched controls for each sample with the CNVkit Python library and command-line software toolkit to infer and visualize copy number from high-throughput DNA sequencing data as described [37]. Copy number calling was performed with default parameters, with the exception of increasing stringency through the use of -m clonal –purity 1 arguments.

### RNA-Seq analysis

Demultiplexed RNA-Seq reads were aligned to Mus musculus mm10 reference genome and expression counts were generated using STAR pipeline on Basepair tech (https://www.basepairtech.com). Gene counts were then normalized, filtered and potential batch effect from tissue was removed using limma v3.46.0 [38] and edgeR v3.32.1 (39). TSNE plots were generated using Rtsne v0.15. Differential gene expression among groups was determined using DESeq2 v1.30.1 [40]. For gene set enrichment analyses (GSEA), pre-ranked gene lists were generated based on shrunken log2 fold changes (LFC) using “ashr” method [41]. GSEA tests were done in clusterProfiler v3.18.1 [42] using KEGG [43] and Reactome [44] database. For differential expression test and GSEA analysis, adjust p-values for false-discovery rate was calculated using the Benjamini-Hochberg method [45]. Plots were generated using ggplot2 v3.3.6.

### High-risk multiple myeloma gene signature analysis

The method for the calculation of EMC-92 signature risk score was modified for use in murine samples from the initial study [18]. Briefly, gene counts of RNA-Seq data were imported as a DESeq object and then applied with DESeq normalization. Variance-stabilizing log2-like transformation of the data was applied using vst command in DESeq package. Human EMC- 92 signature genes were firstly mapped to HGNC symbols (https://www.genenames.org) and then converted to *mus musculus* orthologues. Among all the EMC-92 signature genes, 84 genes were converted to mouse orthologues, in which “C1S”, “FTL”, “DONSON” were found with 2 different orthologues and “SUN1 / GET4” was converted to 2 individual mouse genes. In total, a gene set of 88 genes was used for the calculation. EMC-92 signature risk scores of each sample were then calculated by multiplying the normalized, mean-variance standardized gene expression value with the weighting coefficient factors and the sums of the products (expression value x weighting coefficient) were determined as the EMC-92 signature risk scores.

Analysis of the UAMS-70 gene signature was done through gene set enrichment analysis (GSEA) of the mouse orthologs of the 51 human genes determined to be upregulated in high-risk myeloma patients as described previously [17].

### Mice

CD45.1^+^ transplant recipients were purchased from Jackson Laboratory (stock # 002014) and maintained at Biotron Animal Research Services facility, University of Wisconsin-Madison. Mice were 8-14 weeks old at time of transplant and male and female mice were used in approximately equal proportion. All animal experiments were conducted in accordance with the Guide for the Care and Use of Laboratory Animals and approved by an Animal Care and Use Committee at UW-Madison. The program is accredited by the Association for Assessment and Accreditation of Laboratory Animal Care. All animal experiments in this study are reported in accordance with ARRIVE guidelines (https://arriveguidelines.org).

### Transplantation of myeloma cells

Total splenocytes from a moribund VQ-D2 MM bearing mouse were manually ground through a 70µm filter and washed with sterile PBS containing 2% FBS and antibiotic before being resuspended in 90% FBS/10% DMSO and stored in liquid nitrogen until needed. Frozen VQ- D2 donor cells were gently thawed at 37°C before being washed twice in PBS containing 2% mouse serum (Jackson ImmunoResearch, #015-000-120). Donor cells were then resuspended in 200 μl of PBS containing 2% mouse serum. Eight- to fourteen-week-old CD45.1+ recipient mice were sub-lethally irradiated at 4.0 Gy using a CIX3 irradiator (Xstrahl) and transplanted with 2x10^5^ donor cells via retro-orbital injection.

### Serum protein electrophoresis (SPEP)

Mice were retro-orbitally bled with plain micro hematocrit tubes (Bris, ISO12772). Blood samples were spun in microtainer tubes (BD, 365967) at 4,000x g for 10 minutes to collect serum. Serum was loaded into Hydragel agarose gel (Sebia, 4140) and processed using the Hydrasys instrument (Sebia) following the manufacturer’s instruction. The processed film was scanned, and pixel density of Albumin and γ-globulin bands were quantified using Adobe Photoshop.

### Complete blood count

Peripheral blood samples were collected via retro-orbital bleeding and analyzed with a Hemavet 950FS (Drew Scientific).

### Small compound treatment

For *in vivo* bortezomib treatment, bortezomib (Selleck) was dissolved in sterile PBS and administered at 0.5mg/kg twice a week via intra-peritoneal (IP) injection.

For *in vivo* trametinib treatment, trametinib (Chemietek) was dissolved in 0.5% hydroxypropylmethylcellulose (Sigma) and 0.2% Tween-80 (Sigma) in distilled water (pH 8.0) and administered at 0.25mg/kg via oral gavage daily.

Mice were allocated to treatment groups so that CBC parameters were statistically similar between each group prior to treatment (One-way Analysis of Variance with Tukey-Kramer test). Small compounds were not administered to animals in a blinded manner due to necessary daily preparation of working concentrations for treatment. Animal care staff were blinded to experimental groups during animal assessment. Post-experiment data analysis was not blinded.

### Statistics

For Kaplan–Meier survival curves, survival differences between groups were assessed with the log-rank test, assuming significance at p<0.05. Unpaired, two-way t Test was used to determine significant differences between two groups unless specified. One-way Analysis of Variance with Tukey-Kramer test was used to determine the significance between multiple data sets simultaneously unless specified, assuming significance at p<0.05. Statistical analysis was carried out using GraphPad Prism v9.1.

### Data Availability

All data generated in this study are available upon reasonable request from the corresponding author.

## Results

### B cell receptor repertoire sequencing shows low rates of somatic hypermutation in VQ myeloma

We previously characterized 4-5 VQ myeloma lines, which have distinct sites of MM growth (including the bone marrow, spleen, or lymph node) and dramatically different survival (Figure 1A) [30]. In addition, isotyping of serum antibody from 5 primary VQ mice revealed ubiquitous kappa light chain secretion, with the VQ-D3 line also secreting lambda light chain but no heavy chain (HC) (Figure 1A). All other lines (VQ-D1, D2, D4, and D5) characterized secreted IgA or IgG class-switched HC (Figure 1A), consistent with a post-germinal center derivation [46]. Germinal center B cells undergo somatic hypermutation (SHM) as part of the antibody affinity maturation process [47]. To assess SHM rates in the VQ model, we carried out high-throughput sequencing analysis of BCR repertoire [48] on four primary VQ donor lines (VQ-D1, D2, D4, and D5), as well as two cell lines derived from VQ-D2 recipient mice (4935 and 4938) [30]. Initial library preparation results showed BCR amplicons that were either clonal or oligoclonal with a dominant clone present in the four primary VQ lines (Figure 1B). Sequencing analysis showed monoclonal immunoglobulin heavy chain (IgH) sequences across all primary VQ lines as well as both cell lines (Figure 1C). In terms of immunoglobulin light chain (IgL) sequencing, VQ-D1, D2, and D4 cells showed near clonal IgL, with dominant IgL clones at a frequency of 95% or higher (Figure 1C). Meanwhile, VQ-D5 cells showed oligo-clonal IgL sequences, with three separate clones displaying a frequency of 5% or higher (Figure 1C). Although VQ 4935 and 4938 cell lines maintained clonal IgH similar to parental VQ-D2 cells, both cell lines developed a secondary IgL clone making up approximately 20-25% of sequences (Figure 1C). Interestingly, differences between IgL sequences in VQ cell lines and parental VQ-D2 cells showed clonal selection *in vitro*: the dominant IgL in the VQ 4935 cell line (clone 1) actually arose from a very minor clone in parental VQ-D2 cells with a frequency of only 0.7% (Figure 1C). This same IgL sequence was also present in VQ 4938 cells (clone 2) but did not become the dominant clone.

Subsequent BCR repertoire analysis of all VQ lines showed median SHM rates of IgH chains ≤3.0% (Figure 1D). This SHM frequency is significantly lower than studies in human MM patients, where median SHM rates of ∼8-9% have been previously reported [49, 50]. Corroborating initial library preparation, the VQ 4935 cell line showed a higher mean in SHM (mean = 7.1) compared to other samples, which all had mean SHM of < 4 (Figure 1D). Despite this, median SHM for 4935 cells remained low at 2.64%, while both median and mean SHM rates in VQ 4938 cells were similar as those in parental VQ-D2 cells (Figure 1D). Altogether, BCR repertoire analysis shows that VQ myeloma is characterized by secretion of dominant antibody clones that underwent relatively little affinity maturation prior to BM trafficking.

### Identification of recurrently mutated genes and associated pathways in VQ myeloma cells

Malignant transformation of the VQ model is driven by two common genetic events in human MM: MYC dysregulation and *NRAS Q61R* mutation. However, the prolonged disease latency and phenotypic heterogeneity across different VQ lines suggests the accumulation of additional driver mutations in VQ myeloma cells (Figure 1A). To identify additional mutations, we carried out whole exome sequencing (WES) of CD138+ cells from five VQ mice, as well as corresponding tail samples as germline controls. True mutations were defined as base changes with variant allele frequency (VAF) of >10% in CD138+ cells and <2.5% in matched tail samples. Overall, 11 genes were identified as being recurrently mutated in 2 or more VQ myeloma lines (Figure 2A).

**Figure 2.**
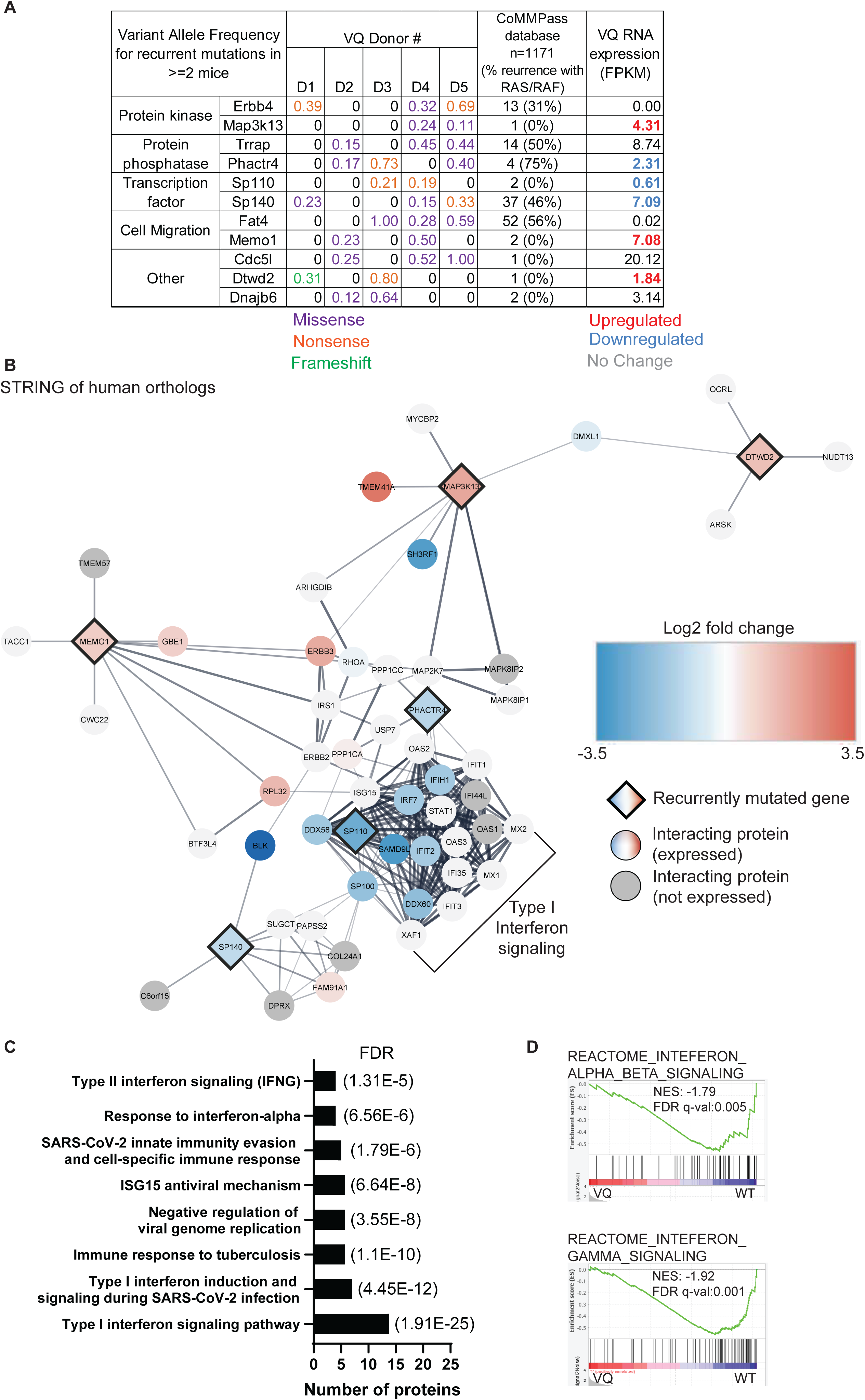
Whole exome sequencing identifies recurrently mutated genes in VQ myeloma cells. Five paired tail DNA (non-leukemia control) and genomic DNA from VQ CD138^+^ B220^-^ plasma cells were subjected to whole exome sequencing as described in Materials and Methods. (A) Recurrently mutated genes (mutated in ≥ 2/4 mice) and their variant allele frequencies (VAF) in VQ cells, frequency of mutation in human orthologs as determined from a cohort of 1,171 MM patient samples from the CoMMPass database, and mRNA expression (as determined by Fragments Per Kilobase of gene per Million mapped reads [FPKM]) in VQ cells are shown. (B) Cytoscape-generated STRING network of recurrently mutated genes with differential RNA expression, as well as 40 closest interacting proteins as determined by STRING analysis. Recurrently mutated genes are represented by diamonds. Interacting proteins are denoted by circles. Genes with differential expression are color-coded. (C) Select list of pathways enriched in genes highlighted in panel B. (D) GSEA of Reactome gene sets for interferon alpha/beta signaling (top) and interferon gamma signaling (bottom) between VQ and control plasma cells. NES, normalized enrichment score. FDR, false discovery rate.

To determine if these genes were recurrently mutated in human MM, mutation status of human orthologs was determined using the Multiple Myeloma Research Foundation (MMRF) CoMMPass database. Of the 11 genes, only *SP140* and *FAT4* were found to be mutated in >3% of patients, Figure 2A), with 37 and 52 out of 1,171 patients, respectively. *SP140* encodes a member of the speckled protein family of chromatin reader proteins [51], which is known to play a role in suppressing Type I interferon signaling in both B cells [52] and macrophages [53]. Frequency of *SP140* mutation ranges from 2.5-7.5% [54,55,56,57] and is found in all stages of MM development from newly diagnosed [54, 55] to drug-refractory [57] disease. *FAT4* encodes a member of the protocadherin family previously identified as putative tumor suppressor gene in the context of breast [58] and colorectal cancers [59]. However, a role of *FAT4* in MM pathogenesis has not been characterized. Frequency of *FAT4* mutation was found in 2-12% of newly diagnosed patients [54, 56], but not significant in rrMM. Of note, *SP140* and *FAT4* mutations are not particularly enriched in MM patients with RAS/RAF pathway mutations as 44% of sequenced patients in the CoMMPass database display mutations in *NRAS, KRAS,* or *BRAF*.

Next, we sought to determine expression of recurrently mutated genes and if their expression levels are altered in myeloma vs control PCs. We generated an expanded RNA-Seq dataset, which includes RNA-Seq analysis of MM cells isolated from multiple tissues from multiple recipient mice in each donor line. Minimum tissue differences were observed in Fragments Per Kilobase of transcript per Million mapped reads (FPKM) from the same mouse. We found that *Erbb4* and *Fat4* were not expressed in PCs (FPKM < 0.1) and 6 of the 11 recurrently mutated genes were differentially expressed in BM VQ cells compared to control PCs (Figure 2A, far right column). Among them, *Map3k13*, *Memo1*, and *Dtwd2* were upregulated, while *Phactr4*, *Sp110*, and *Sp140* were downregulated in VQ myeloma cells. Interestingly, none of these genes have previously been characterized in the context of MM. To determine the potential pathways affected by their dysregulation in the VQ model, we used the Cytoscape network analysis package [60] to create a protein interaction map using human orthologs of the six recurrently mutated and dysregulated genes, along with their 50 closest interacting proteins as determined by Search Tool for the Retrieval of Interacting Genes/Proteins (STRING) analysis [61] (Figure 2B). We further integrated our RNA-Seq data with the protein-protein interaction network (see Figure 2B legend at bottom right). Intriguingly, we noticed that most of the genes in this interaction network were downregulated at the transcriptional level in VQ cells (Figure 2B). STRING functional enrichment of these proteins showed a significant association with both Type I and Type II Interferon signaling, as well as immune responses to viral and tuberculosis infection (Figure 2C). Corroborating these findings, gene set enrichment analysis (GSEA) of VQ MM cells showed downregulation of both Type I and Type II interferon signaling compared to control PCs (Figure 2D). Altogether, our data suggests that interferon signaling is downregulated in VQ myeloma cells.

### Copy number variation analysis stratifies VQ myeloma lines into two clusters based on recurrent amplification of chromosome 3 and monosomy of chromosome 5

CNVs are frequent in MM and play a well-established role in predicting patient prognosis [62]. Correspondingly, genetic characterization of the 5T and Vĸ*MYC transgenic mouse models also show a prevalence of CNVs in murine myeloma cells [27, 28]. We thus sought to carry out CNV analysis of the five VQ lines using our WES data. Amplification (amp) of chromosome (chr) 3 was observed in three of the five VQ lines (Cluster I: VQ-D2, VQ-D3, and VQ-D5) (Figure 3A), while monosomy of chr5 was observed in the other two lines (Cluster II: VQ-D1 and VQ-D4, Figure 3B). In addition, monosomy chr8, trisomy chr7, and trisomy chr15 were observed in VQ- D1, VQ-D3, and VQ-D4, respectively (Figure 3A-B). Similar CNVs have also been observed in other transgenic MM models: the 5T2 line of the 5T model shows both trisomy 3 and monosomy 5 [27], whereas monosomy 5 is observed in approximately 50% of primary Vĸ*MYC lines [28].

**Figure 3.**
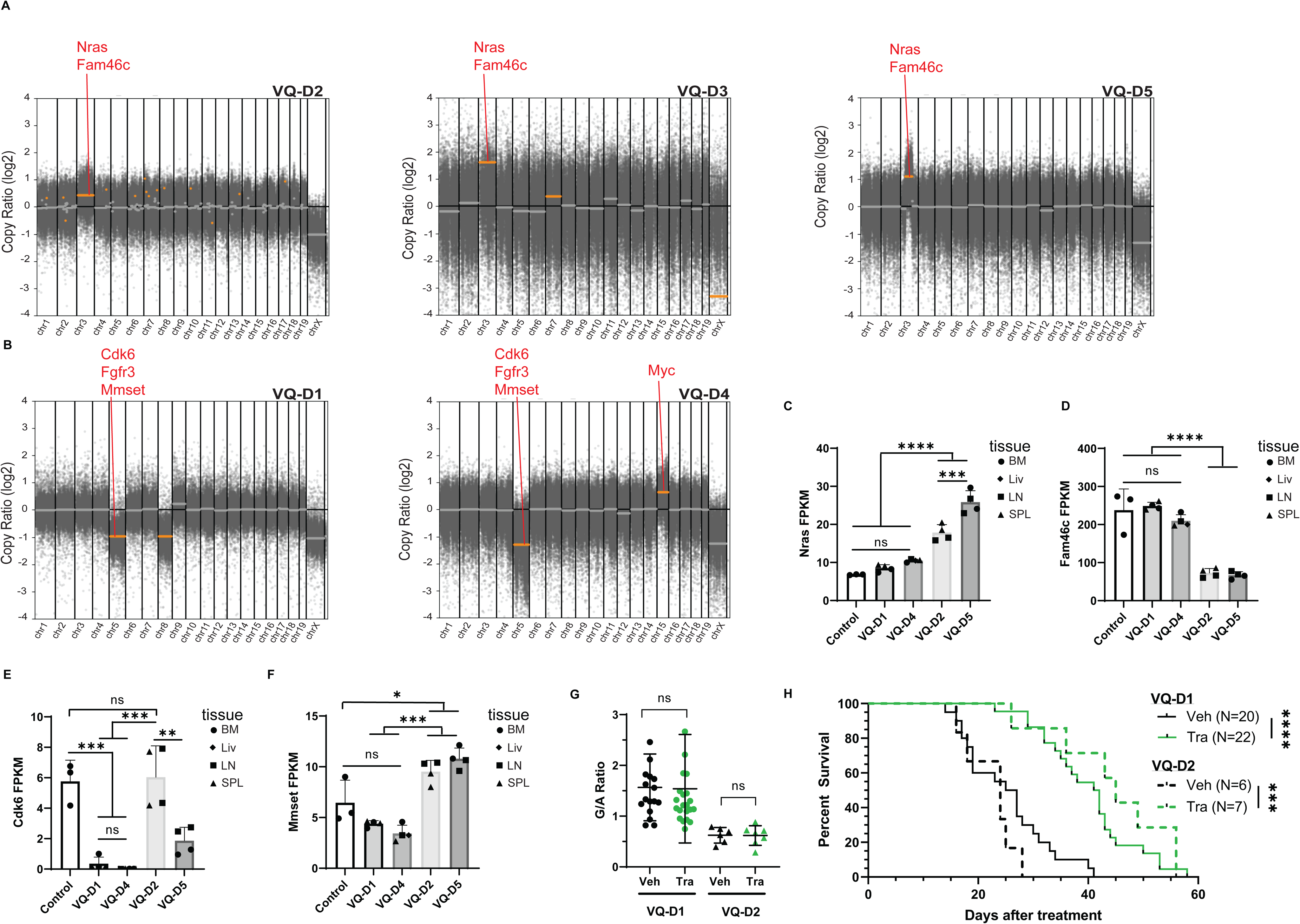
Copy number variation (CNV) analysis stratifies VQ cells based on recurrent amplification of chromosome 3 and monosomy chromosome 5. (A-B) CNV analysis was performed using the whole exome sequencing data as described in Materials and Methods. Orange dots indicate significant changes in log2 copy ratio for a given call segment in plasma cells compared to non-leukemia control samples. Location and name of tumor suppressors and oncogenes related to myeloma pathogenesis are shown in red. CNV plots are grouped according to recurrent CNV status. (C-F) Transcript levels of genes highlighted in red in panels A and B are shown in CD138+ B220- cells from control and VQ recipient mice. FPKM, Fragments Per Kilobase of transcript per Million mapped reads. One-way analysis of variance with Tukey’s post-test was performed. Tissue of origin for individual samples is denoted by legend. (G-H) CD45.1 recipient mice were sub-lethally irradiated and injected with bone marrow cells from moribund VQ-D1 donor mouse or splenocytes from moribund VQ-D2 donor mouse. Six weeks (VQ-D1) or two weeks (VQ-D2) post-transplant, mice were treated with vehicle or trametinib. (G) Serum protein electrophoresis was performed to quantify the γ-globulin/Albumin (G/A) ratios in VQ-D1 and VQ-D2 recipient mice at day 21 of treatment. Two-sided t-test was performed. (H) Kaplan-Meier survival curves were plotted against days after treatment. Log-rank test was performed. Note: VQ-D1 results are combined from historical [30, 31] and new data. ns, not significant. *, p<0.05. **, p<0.01. ***, p<0.001. ****, p<0.0001.

CNVs identified in the VQ model include several tumor suppressor genes and oncogenes that are highly relevant for MM pathogenesis (Figure 3A-B, highlighted in red). Notably, chr3 includes the proto-oncogene *Nras*, as well as *Fam46c* encoding a non-canonical poly(A) polymerase that is frequently mutated in MM and acts as a tumor suppressor for MM development [63, 64]. Chr5 includes *Fgfr3* and *Mmset*, both of which are overexpressed in t(4;14) MM patients [12], and cyclin-dependent kinase 6 (*Cdk6*) that was recently found to be over-expressed in immunomodulatory drug-resistant MM cells [65]. Finally, chr15 includes the transcription factor *Myc*, frequently dysregulated in MM [66].

We next wanted to determine if CNVs of the affected genes correlate to their transcriptional changes in VQ vs control PCs. We previously reported moderate overexpression of Myc in all VQ MM lines due to the Vk*MYC transgene [30] and absence of Fgfr3 expression in both control and VQ myeloma PCs [31]. Using the expanded RNA-Seq dataset, we observed an approximately two-fold increase in *Nras* expression in Cluster I (VQ-D2 and VQ-D5) cells with chr3 amplification compared to control and Cluster II VQ lines without chr3 amplification (Figure 3C). Despite chr3 amplification, expression of *Fam46c* was significantly lower in Cluster I cells compared to Cluster II and control PCs, suggesting an epigenetic mechanism involved in *Fam46c* downregulation (Figure 3D). In Cluster II (VQ-D1 and VQ-D4) cells with monosomy chr5, *Cdk6* was nearly undetectable (average FPKM < 0.5), whereas VQ-D5 cells had a 3-fold decrease compared to VQ-D2 cells and control PCs (Figure 3E). Despite monosomy chr5, *Mmset* expression in Cluster II MM cells was comparable to that in control PCs, whereas it was approximately two-fold higher in Cluster I cells (Figure 3F). Taken together, our data demonstrate that gene copy numbers do not necessarily correlate with mRNA levels in VQ cells, suggesting that additional epigenetic mechanisms play an important role in controlling gene expression.

We previously showed that daily treatment of trametinib (Tra), a potent MEK inhibitor, did not lower myeloma disease burden in VQ-D1 recipients as measured by the ratio of serum gamma-globulin to albumin (G/A) using serum protein electrophoresis (SPEP) (Figure 3G). However, it significantly prolonged the survival of diseased mice [30, 31] (Figure 3H). Because Cluster I VQ cells show increased *Nras* copy number and mRNA expression, we wanted to determine if VQ- D2 cells were resistant to MEK inhibition. Here, we followed the same treatment schemes in VQ-D2 recipient mice as previously carried out in VQ-D1 mice (see Materials and Methods). Two weeks after VQ-D2 transplantation, recipients were divided into two groups with comparable CBC parameters and treated with vehicle (Veh) or Tra. Three weeks later, treatment efficacy was assessed via G/A ratio using SPEP assay. Despite the amplification of chr3 with the *Nras* locus in Cluster I myeloma lines, daily Tra treatment provided a survival benefit to VQ-D2 recipients similar to what has been previously observed in VQ-D1 mice (Figure 3G and 3H).

### Gene transcription-based clustering of VQ lines yields highly consistent result with the CNV study

Our CNV analysis separated VQ lines into two distinct clusters based on their characteristic genomic changes, which may cause consistent global transcriptional changes in VQ cells. To test this idea, we performed unsupervised hierarchical clustering analysis of all control and VQ samples based on the transcriptional profiles generated in RNA-Seq. Not surprisingly, these samples split into three groups: Cluster I with chr3 amplification (VQ-D2 and VQ-D5), Cluster II with monosomy chr5 (VQ-D1 and VQ-D4), and control PCs (Figure 4A). Further analysis of sample similarity via t-distributed stochastic neighbor embedding (tSNE) (Figure 4B) and principal component analysis (PCA) (Figure S1) confirmed that VQ myeloma lines cluster into two distinct transcriptional subtypes. This result was not affected by tissue origins of MM cells as similar clustering result was obtained when tissue effect was removed (Figure S2). More importantly, the transcriptional clusters mirror recurrent CNVs observed in these VQ lines.

**Figure 4.**
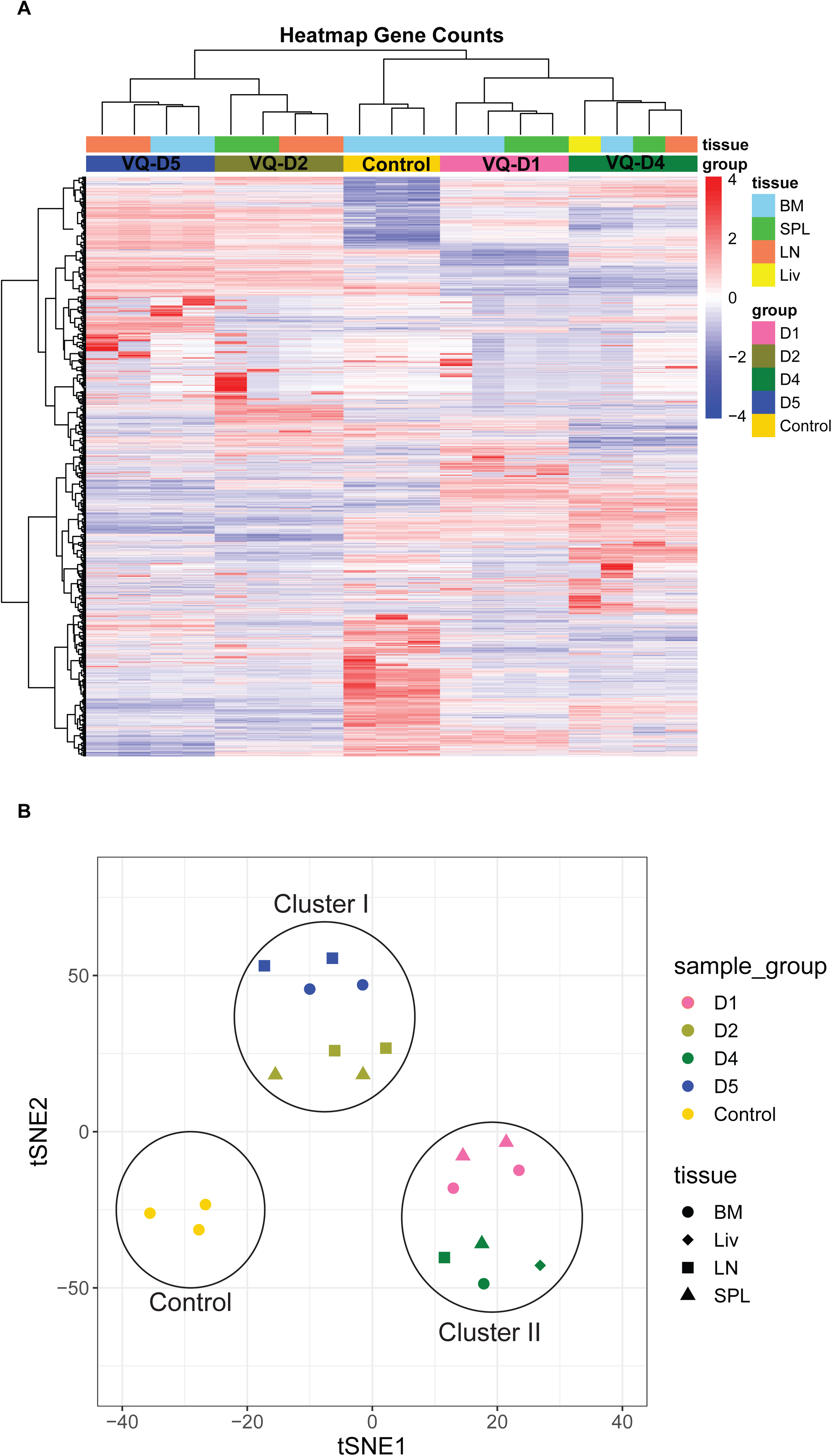
RNA-Seq analysis reveals two distinct transcriptional clusters of VQ myeloma. Bulk RNA-Seq analysis was performed using flow sorted CD138+ B220- CD45.2+ cells from bone marrow (BM) of control mice (n=3) and BM, spleen (SPL), lymph node (LN), or liver (Liv) of multiple VQ-D1, VQ-D2, VQ-D4, and VQ-D5 recipients. (A) Clustered heat map of RNA-seq gene count data. Samples are color-coded by tissue sites and VQ lines as indicated. (B) tSNE analysis of gene counts data of VQ myeloma samples. Samples are color-coded by VQ lines as in panel A. Tissue of origin for individual samples is denoted by legend.

### Cluster I VQ cells display upregulation of growth pathways and high-risk myeloma gene signatures

We next sought to differentiate transcriptional activity between Cluster I and Cluster II VQ myeloma. GSEA of hallmark signaling pathways, oncogenic signatures, and relevant MM-related gene signatures showed numerous pathways significantly upregulated in Cluster I VQ cells, but relatively few in Cluster II (Figure 5A). Interestingly, we observed that essential cancer growth pathways were highly upregulated in Cluster I VQ cells (Figure 5A), including E2F targets (Figure 5B), G2M checkpoint (Figure 5C), and Myc target pathways (Figure 5D). Moreover, Cluster I VQ cells showed significant upregulation of the UAMS-70 and EMC-92 high-risk MM gene signatures compared to Cluster II VQ lines (Figures 5E). Similar results were observed when we compared hrMM gene signatures of VQ clusters to the transplantable Vĸ*Myc line t-Vk12653 [67]. While the UAMS-70 gene signature was enriched in Cluster I VQ compared to t-Vk12653 (Figure S3A), neither of the hrMM gene signatures was enriched in VQ Cluster II cells (Figure S3B). Because EMC92 gene signature includes both positive and negative risk genes with different weights, we calculated the EMC-92 risk scores in Cluster I and II as well as t-Vk12653 MM cells based on a modified version of the published algorithm [18]. The EMC-92 risk score of Cluster I MM cells was significantly higher than those of Cluster II and t-Vk12653 cells (Figure S3C). These analyses clearly defined VQ Cluster I myeloma lines as having high-risk gene expression in comparison to either VQ Cluster II or t-Vk12653 cells.

**Figure 5.**
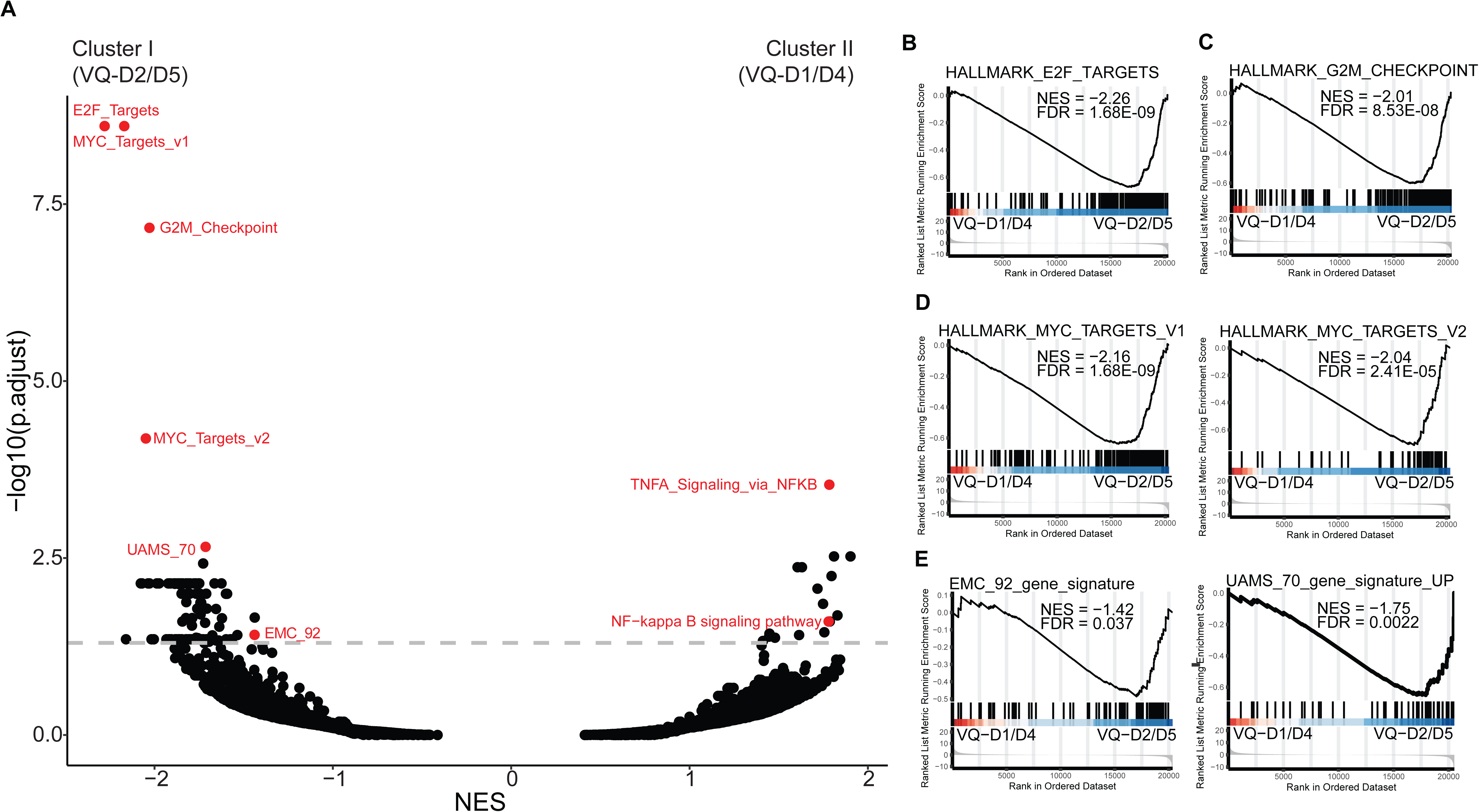
VQ Cluster I myeloma cells have increased expression of Hallmark growth pathways and high-risk MM gene signatures. (A) Overview of gene set enrichment analysis between Cluster I (VQ-D2/D5) and Cluster II (VQ-D1/D4) VQ myeloma cells. Relevant pathways are highlighted in red. (B-E) GSEA plots comparing Cluster I to Cluster II for (B) Hallmark_E2F_Targets, (C) Hallmark_G2M Targets, (D) Hallmark_MYC_Targets_V1 (left) Hallmark_MYC_Targets_V2 (right), and (E) EMC-92 hrMM gene signature (left) and UAMS-70 hrMM gene signature (right). NES, normalized enrichment score; FDR, false discovery rate; p. adj., adjusted P-value.

One of the most significantly upregulated genes in Cluster II VQ is the transcription factor Pbx1, which has recently been characterized as a driver oncogene in the context of chr1q-amp hrMM [68]. PBX1, along with its downstream target FOXM1, were found to drive proliferation and drug resistance in chr1q-amp MM patients [68]. Interestingly, despite loss of *Pbx1* in Cluster I VQ (Figure S4A), transcriptional levels of *Foxm1* in Cluster I VQ cells were significantly higher than that in Cluster II VQ cells or control PCs (Figure S4B). We further assessed the PBX1-FOXM1 axis in VQ myeloma using the recently characterized PBX1 gene signature [68] and a previously established FOXM1 pathway signature [69]. Interestingly, we found that while neither VQ-D2 nor VQ-D5 cells expressed detectable PBX1, Cluster I VQ was enriched for both the PBX1 (Figure S4C) and FOXM1 (Figure S4D) gene signatures compared to both Cluster II VQ and control PCs. In addition, despite high *Pbx1* mRNA expression in VQ-D1 and VQ-D4 cells, neither gene signature was significantly enriched in Cluster II VQ compared to control PCs. The enrichment of PBX1 and FOXM1 gene signatures in Cluster I VQ cells is consistent with their high-risk nature.

### Cluster I and Cluster II VQ show distinct responses to bortezomib *in vivo*

When studying the pathways enriched in Cluster II vs Cluster I VQ MM cells, we found that both NF-ĸB and TNF-α signaling via NF-ĸB pathways were significantly upregulated in Cluster II VQ (Figure 6A and 6B). We also observed comparable expression of *Il2rg* and increased expression of *Cd74* and *Tnfaip3* in Cluster II vs Cluster I myeloma cells (Figure S5A-C). The expression levels of these three genes are positively correlated to bortezomib (Btz) response in human MM [70]. In addition, in comparison to control PCs, Cluster I myeloma cells overexpressed *Psmb1*, *Psmb2*, and *Psmb 5* (Figure 6E), which encode the potential Btz targeted proteasome subunits [71]. By contrast, Cluster II myeloma cells only overexpressed *Psmb1* and *Psmb 5* (Figure 6E). Together, our results suggest that Cluster I and II myeloma cells may respond differently to Btz treatment.

**Figure 6.**
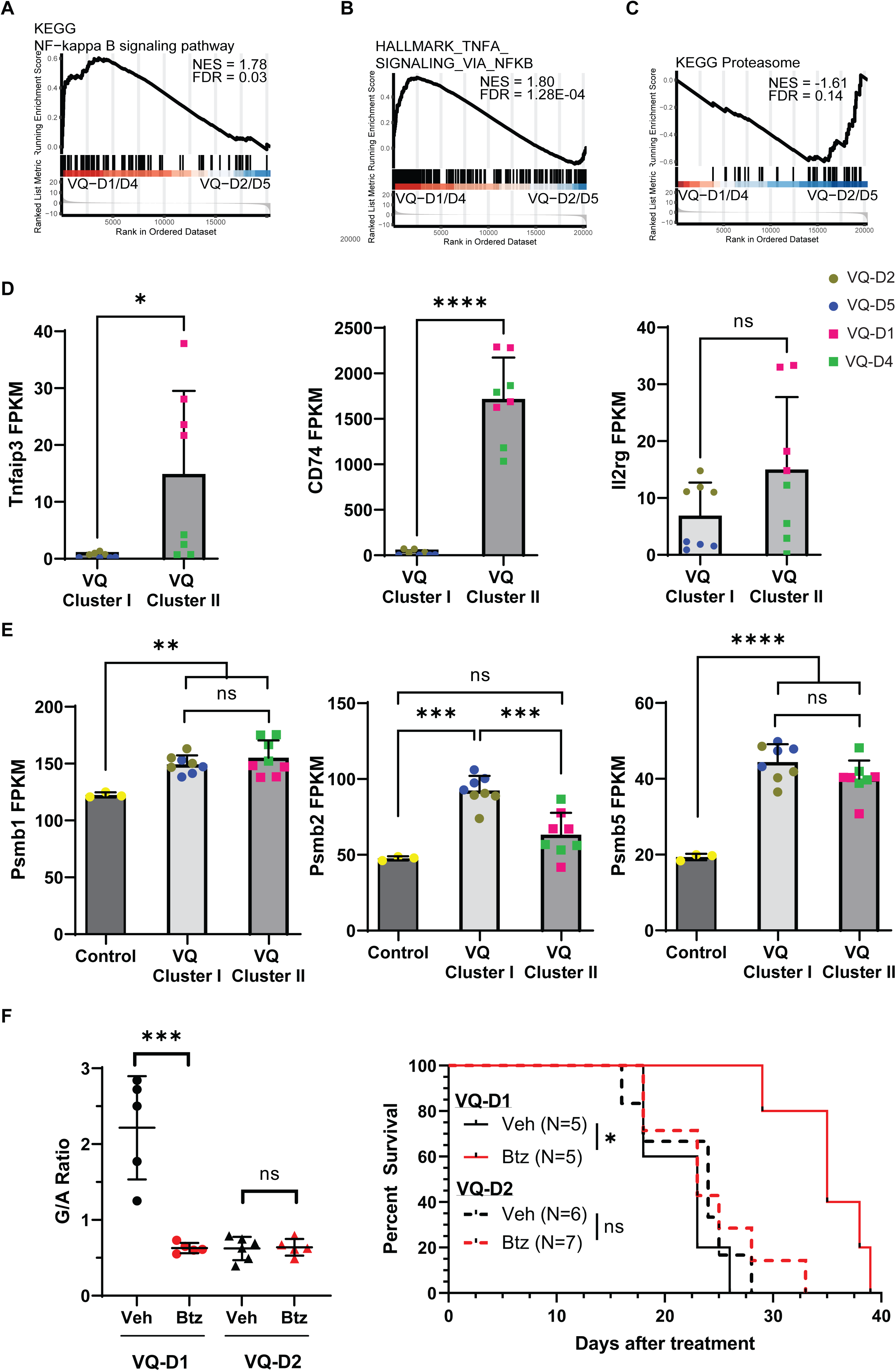
Cluster I and Cluster II VQ cells display distinct responses to bortezomib *in vivo.* (A-C) Gene set enrichment analysis of proteasome inhibitor related pathways between Cluster I (VQ-D2/D5) and Cluster II (VQ-D1/D4) VQ myeloma cells. KEGG_ NF-kappa B signaling pathway (A), Hallmark_TNFA signaling via NFKB (B), and KEGG_Proteasome pathway (C). (D) Transcript levels of NF-ĸB related genes *Tnfaip3*, *CD74*, and *Il2rg* in CD138+ B220- cells from Cluster I and II VQ myeloma mice. Two-sided t-Test was performed. VQ donor of origin is color-coded as indicated. Results are presented as mean + SD. FPKM, Fragments Per Kilobase of transcript per Million mapped reads. (E) Transcript levels of proteasome related genes *Psmb1*, *Psmb2*, and *Psmb5* in CD138+ B220- cells from control plasma cells (Control) and Cluster I and II VQ cells. Two-sided t-Test was performed. VQ donor of origin is color-coded as indicated. Results are presented as mean + SD. (F-G) CD45.1 recipient mice were sub-lethally irradiated and injected with bone marrow cells from moribund VQ-D1 donor mouse or splenocytes from moribund VQ-D2 donor mouse. Six weeks (VQ-D1) or two weeks (VQ-D2) posttransplant, mice were treated with vehicle or bortezomib as described in Materials and Methods. (F) Serum protein electrophoresis was performed to quantify the γ-globulin/Albumin (G/A) ratios in VQ-D2 recipient mice at day 21 of treatment. Two-sided t-Test was performed. (G) Kaplan-Meier survival curves were plotted against days after treatment. Log-rank test was performed. Note: VQ-D1 results are taken from historical data [31]. FDR, false discovery rate; NES, normalized enrichment score; ns, not significant; p. adj., adjusted P-value. *, p <0.05; **, p <0.01; ***, p <0.001; ****, p <0.0001.

We previously established the single-agent efficacy of inhibiting the proteasome pathway in VQ- D1 via Btz [31]. Here, we followed the same Btz treatment scheme in VQ-D2 recipient mice as previously established in VQ-D1 (see Materials and Methods). Two weeks after VQ-D2 transplantation, recipients were divided into two groups with comparable CBC parameters and treated with vehicle (Veh) or Btz. Three weeks later, treatment efficacy was assessed via G/A ratio. In contrast to VQ-D1 mice, in which Btz lowered myeloma burden after three weeks of treatment and provided a moderate but significant survival boost [31], this same Btz treatment showed no efficacy in VQ-D2 mice and did not provide any survival benefit compared to Veh treated mice (Figure 6F). This *in vivo* study validated our molecular characterization, demonstrating that Cluster I VQ-D2 has *de novo* Btz resistance and can thus be considered higher risk than Cluster II VQ-D1.

## Discussion

Previously, our group developed the Ras-driven VQ myeloma model and phenotypically characterized five lines derived from primary VQ mice (Figure 1A) [30]. In the current study, we combined BCR repertoire sequencing, WES, and CNV with RNA-Seq to characterize the genomic and transcriptomic landscapes of VQ MM lines. Both genomic and transcriptional analyses stratified VQ lines into two distinct clusters. Cluster I includes VQ-D2 and VQ-D5, which carry chr3 amplification and display both UAMS-70 and EMC-92 hrMM gene signatures vs Cluster II VQ and t-Vk12653 Vĸ*Myc myeloma. By contrast, Cluster II includes VQ-D1 and VQ-D4, which harbor monosomy chr5 and express hrMM gene signatures comparable to t- Vk12653 Vĸ*MYC myeloma. Consistent with their molecular classification, Cluster I myeloma cells showed de novo resistance to Btz treatment *in vivo*, while Cluster II myeloma cells exhibited a reduced response to Btz. Interestingly, both Cluster I and II VQ lines responded to Tra. Our molecular stratification of VQ lines provide a foundation to predict the potential outcomes of therapeutic regiments in different populations of human MM patients.

A negative association between the rate of SHM and patient prognosis has been previously established in CLL [72]. However, such an association is unclear in the context of MM, as SHM has been found to have a positive correlation with patient prognosis [49] or none at all [50]. With the exception of the VQ 4935 cell line, BCR repertoire analysis showed relatively low rates of SHM in VQ cells compared to human myeloma [49], with mutation frequencies ≤3.0%. Although median SHM frequency was slightly higher in Cluster I VQ (VQ-D2/D5) (Figure 1B), there was no significant correlation with median survival of different VQ lines (Figure 1A). Notably, SHM in VQ cells is comparable to levels in primary Vĸ*MYC mice, where a median frequency of 2.6% in IgH was observed [24].

WES of five VQ lines identified 11 recurrently mutated genes (Figure 2). Human orthologs of two of these genes, *FAT4* and *SP140*, have been identified as recurrently mutated in numerous MM patient sequencing studies [36–40]. Combining mRNA expression and mutation frequencies during human MM development, we believe that *FAT4* and *SP140* mutations play distinct roles in MM development. *Fat4* expression was nearly undetectable at the mRNA level in both VQ and control PCs (Figure 2A). RNA-Seq data compiled by the MMRF’s CoMMPass database shows similarly low expression, with 80% of patients having <2 transcripts per million reads for the *FAT4* gene (data not shown). Moreover, although *FAT4* mutations were detected in a fraction of newly diagnosed patients [54, 55], they remained stable in a recent temporal sequencing study of 62 MM patients [73]. Together, these data suggest that *FAT4* mutations serve a passenger role in myeloma progression. By contrast, *Sp140* was found to be expressed in both control and VQ PCs and its mRNA level was significantly downregulated in VQ MM cells (Figure 2A). In MM patients, frequency of *SP140* mutation increases in more advanced stages of disease [54,55,56,57] and also increased over time in the same temporal sequencing study [73]. Altogether, these data suggest that *SP140* mutation plays an important role in MM development and progression.

*SP140* encodes a member of the speckled protein family of chromatin reader proteins [51], and has been previously characterized for its role in downregulating interferon signaling in select immune cells [52, 53]. *Sp110*, encoding a homolog of Sp140 [51], was also found to be recurrently mutated in VQ cells and downregulated at the transcriptional level compared to control PCs (Figure 2A). STRING analysis showed that Sp110, along with Phosphatase and actin regulator 4 (Phactr4), whose gene was also recurrently mutated in VQ MM cells, share interactions with several proteins that were also downregulated at the mRNA level in VQ cells (Figure 2B). Pathway analysis of this group of proteins found that they were enriched in Type I and Type II interferon signaling (Figure 2C). Although Phactr4 has been found to act as a tumor suppressor in hepatocellular carcinoma due to its inhibition of the IL-6/STAT3 pathway [74] and both Sp110 and Sp140 were previously implicated in interferon signaling in response to tuberculosis infection [75], none of these genes have previously been implicated in interferon signaling in the context of MM. Evolutionarily, attenuating Type I and Type II interferon signaling could be beneficial to MM immune evasion and survival. Prior to the development of current novel therapy regiments, IFN-α2b was used as a MM treatment [76] with limited long-term use due to its systemic toxicity [77]. More recent studies in which IFN-α2b is conjugated to antibodies targeting CD38 [78] or HLA-DR [79] on the myeloma cell surface have also shown both *in vitro* and *in vivo* efficacy. Therefore, the role of Sp110 and Sp140 in shaping the interferon response in VQ as well as in human MM is an interesting question worth future consideration.

Large-scale chromosomal changes, including translocations and aneuploidies, have significant impacts on treatment outcomes in MM patient prognosis. For example, use of the BCL-2 inhibitor venetoclax for t(11;14) MM [80] or the innate resistance of t(14;16) MM to PIs due to c-MAF overexpression [81, 82] are well documented. Therefore, we performed CNV analysis in five VQ lines and identified two recurrent, mutually exclusive events: amplification of chr3, which was present in Cluster I (VQ-D2, D3, and D5; Figure 3A), and monosomy chr5, which was present in Cluster II (VQ-D1 and D4; Figure 3B). Neither of these CNVs are unique to the VQ model. Full gain(chr3) has been identified in the 5T2 line, while the 5TGM1 cells showed partial duplication [27]. Interestingly, monosomy 5 is a unifying CNV across all three murine MM models. Monosomy 5 was also observed in 5T2 cells, while only partially deleted in 5T33vv and 5TGM1 lines. Noticeably, monosomy 5 was present in 13/26 sequenced Vĸ*MYC lines [28]. The high incidence of monosomy chr5 across multiple MM models warrants further investigation.

Consistent with our CNV analysis, gene transcriptional analyses of RNA-Seq data, including non-supervised hierarchal clustering, PCA, and tSNE, identified two distinct transcriptional clusters (Figure 4 and Figure S1) regardless of their discrete tissue origins (Figure S2).

Extensive pathway analysis between these two clusters as well as in comparison to the t- Vk12653 Vk*MYC line revealed the hrMM order: Cluster I > Cluster II = t-Vk12653 (Figure 5 and S4), On the other hand, Cluster II (VQ-D1/D4) cells were enriched for the TNF-α/NF-ĸB signaling (Figure 6A and 6B) and increased expression of genes linked to Btz response in patients (Figure 6C). These transcriptional differences led us to speculate that Cluster I cells may be more resistant to PIs than Cluster II cells. In our initial study, treatment of VQ-D1 recipient mice with Btz following a dosing schedule found to be effective in the Vĸ*MYC model (1.0mg/kg Btz treatment on days 1,4,8,11) [67] was not effective as a single agent, but did provide a survival benefit when combined with the MEK inhibitor selumetinib [30]. However, in a more recent study, we found that reducing Btz dosage to current clinical practices provided a modest but significant boost to VQ-D1 survival and significantly reduced G/A ratio after three weeks of treatment [31] (Figure 6F). By contrast, in our current study we found that the same Btz treatment was completely ineffective against VQ-D2 cells (Figure 6F), corroborating our molecular analysis.

We observed that despite chr3 amplification and Nras overexpression in Cluster I MM, both VQ- D1 and VQ-D2 MM cells responded to Tra treatment (Figure 3). By contrast, 5T2 MM cells showed increased resistance to Tra compared to other 5T lines, though this is complicated by the fact that the genomic region containing *Kras* is also duplicated in 5T2 cells [27].

Nonetheless, our results provide a strong rationale to develop Tra-based combination therapies in hrMM and rrMM, particularly in the context of RAS mutations.

Altogether, our data further elucidate the genomic and transcriptional landscapes of the VQ model. These molecular characterizations, along with functional validation via *in vivo* treatment experiments, supports an intra-model stratification where Cluster II VQ (VQ-D1/D4) models Ras-driven MM with some hrMM features comparable to non-Ras-driven MM (e.g. t-Vk*MYC), whereas Cluster I VQ (VQ-D2/D5) represents ultra-high risk multiple myeloma. Future studies using the VQ model as pre-clinical platform must necessarily take these intra-model differences into account when designing mechanistic studies or testing novel treatments.

## Acknowledgments

We would like to thank the University of Wisconsin Carbone Comprehensive Cancer Center (UWCCC) for use of its Shared Services (Small Molecule Screening Facility, Flow Cytometry Laboratory, Transgenic Animal Facility, and Experimental Pathology Laboratory) to complete this research. We would also like to thank Dr. Robert Burns for his assistance in initiating the CNV study. This work was supported by a postdoctoral fellowship from the NIH grant T32 GM081061 to EF, the startup fund 252840-00 from Marshfield Clinic Research Foundation to ZW, R01CA212413 to AR, R01CA252937 to FA, R01CA152108 to JZ, R01AI079087 and R01HL130724 to DW, and additional support from the Trillium Fund, UWCCC Developmental Therapeutics Program Pilot Awards, and Immunotherapy Pilot Award.

