## Supplemental Figure Legends for "Genomic and transcriptional profiling stratifies VQ myeloma lines into two clusters with distinct risk signatures and drug responses"

**Supplemental Information**

**Supplemental Figure Legends**

**Figure S1. Principal component analysis of RNA-seq data show VQ cells group into two distinct transcriptional clusters.** Bulk RNA-Seq analysis was performed as described in Figure 4. PCA plot of gene count clustering analysis. Samples are color-coded by group as indicated. Tissues of origin for individual samples are denoted by legend.

**Figure S2. Tissue site of origin does not affect unsupervised clustering of VQ myeloma and wildtype plasma cells.** Bulk RNA-Seq analysis was performed with tissue effect taken into account as described in Figure 4. (A) Clustered heat map of RNA-seq gene count data. Samples are color-coded by tissue sites and VQ lines as indicated. Tissue effect of individual samples was removed as described in Materials and Methods. (B) tSNE plot of gene count clustering analysis. Samples are color-coded by VQ lines as in panel A. Tissues of origin for individual samples are denoted by legend. (C) PCA plot of gene count clustering analysis. Samples are color-coded by VQ lines as in panel A. Tissues of origin for individual samples are denoted by legend.

**Figure S3. High-risk multiple myeloma gene signatures are enriched in VQ Cluster I compared to VQ Cluster II or t-Vk12653 Vĸ*Myc cells.** GSEA plots comparing the UAMS-70 and EMC-92 high-risk multiple myeloma gene signatures between Cluster I VQ cells (VQ-D2/D5) and t-Vk12653 cells (A) and between Cluster II VQ cells (VQ-D1/D4) and t-Vk12653 cells (B). (C) EMC-92 risk scores calculated using clinical risk algorithm (see Materials and Methods) for control plasma cells, Cluster I and II VQ cells, and t- Vk12653 cells. Two-sided t-test with Holm Bonferroni Correction was performed. Samples are color-coded and tissues of origin for individual samples are denoted by legend. FDR, false discovery rate; NES, normalized enrichment score. **, p <0.01; ***, p, <0.001.

**Figure S4. Cluster I VQ lines are enriched for the high-risk PBX1 and FOXM gene signatures.** (A, B) Transcript levels of transcription factors Pbx1 (A) and Foxm1 (B) in CD138^+^ B220^-^ cells from control or VQ recipient mice. Two-sided t-Test was performed. VQ donor of origin is color-coded as indicated. Results are presented as mean + SD. (C) GSEA plots comparing the PBX1 signature between VQ-D1/D4 and VQ-D2/D5 (top), control and VQ-D1/D4 (middle), and control and VQ-D2/D5 (bottom). (D) GSEA plots comparing the FOXM1 pathway between VQ-D1/D4 and VQ-D2/D5 (top), control and VQ-D1/D4 (middle), and control and VQ-D2/D5 (bottom). FDR, false discovery rate; FPKM, Fragments Per Kilobase of transcript per Million mapped reads; NES, normalized enrichment score. **, p <0.01; ****, p, <0.0001.
