## Supplementary figures and images for "Genomic and transcriptional profiling stratifies VQ myeloma lines into two clusters with distinct risk signatures and drug responses"

### Supplemental Figure 1

**Figure S1-Zhang**

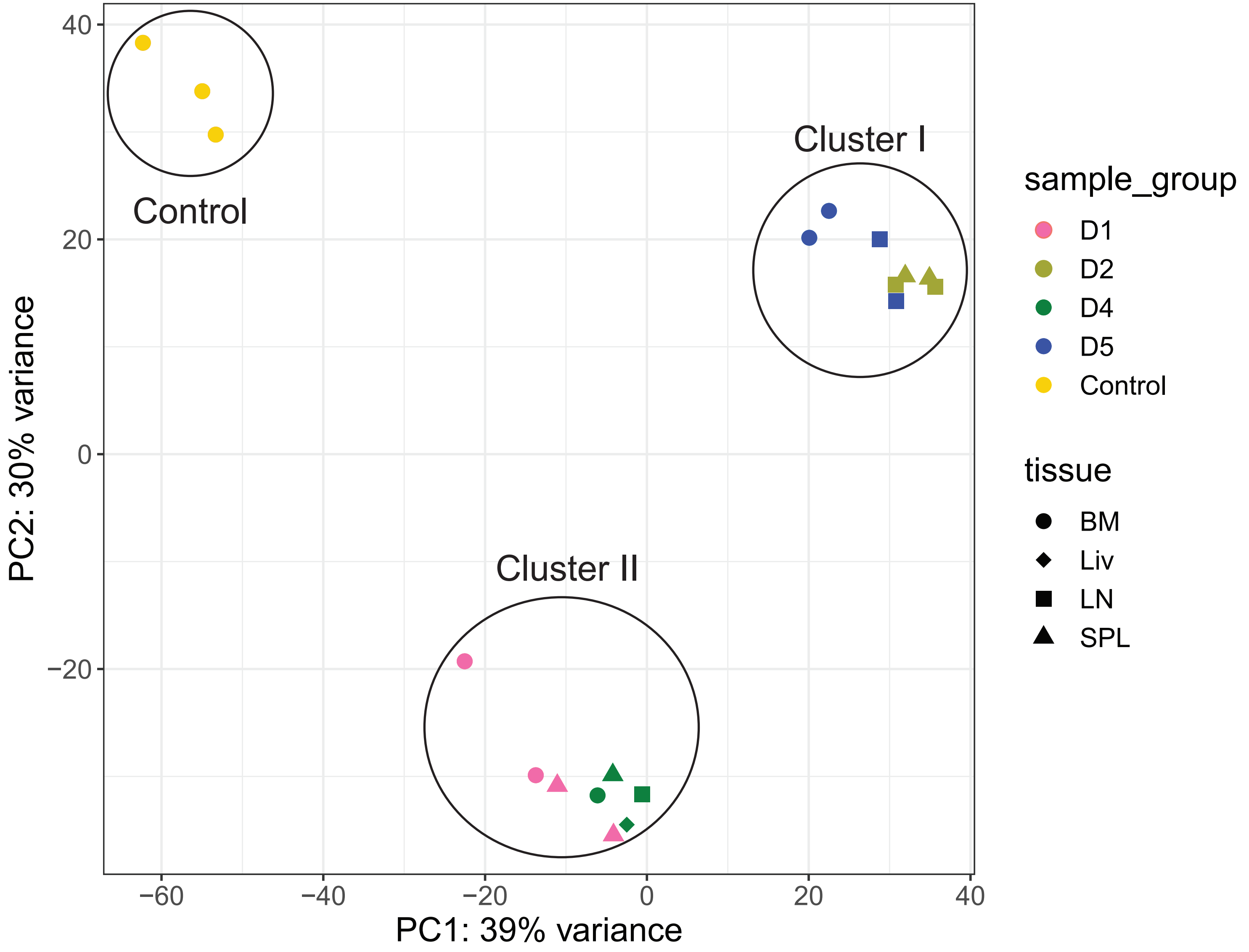

### Supplemental Figure 2

Figure S2-Zhang

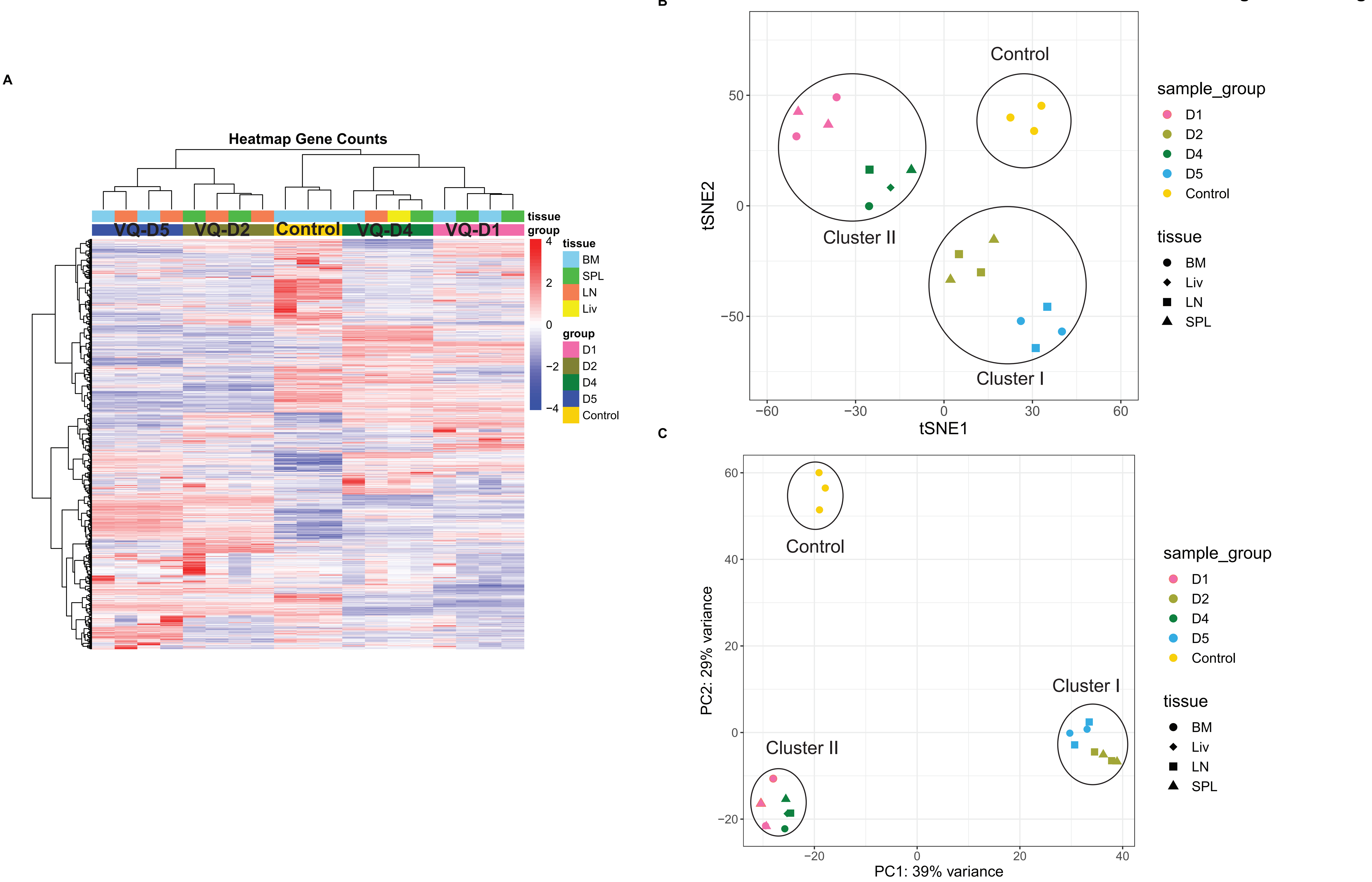

### Supplemental Figure 3

**Figure S3-Zhang**

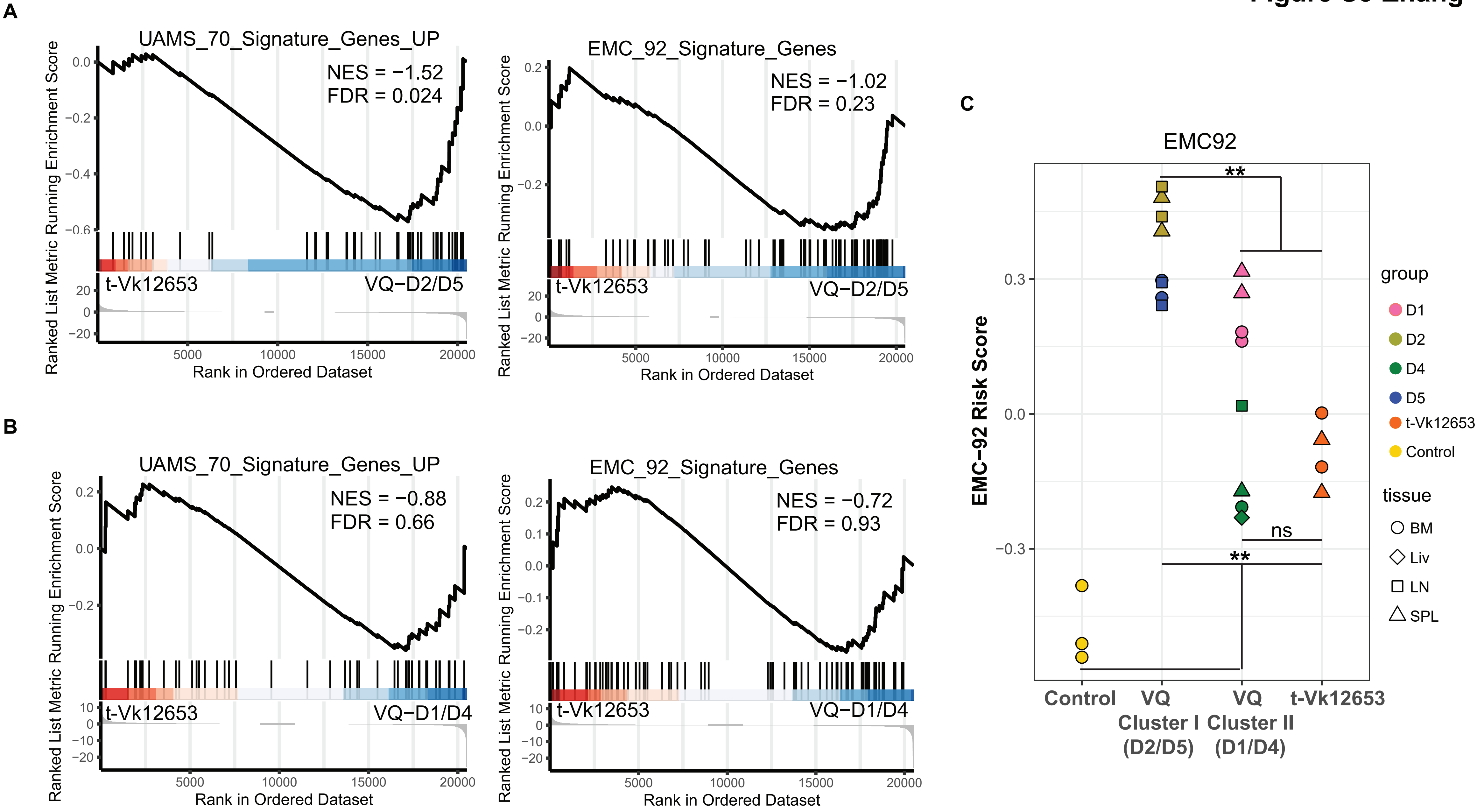

### Supplemental Figure 4

**Figure S4-Zhang**

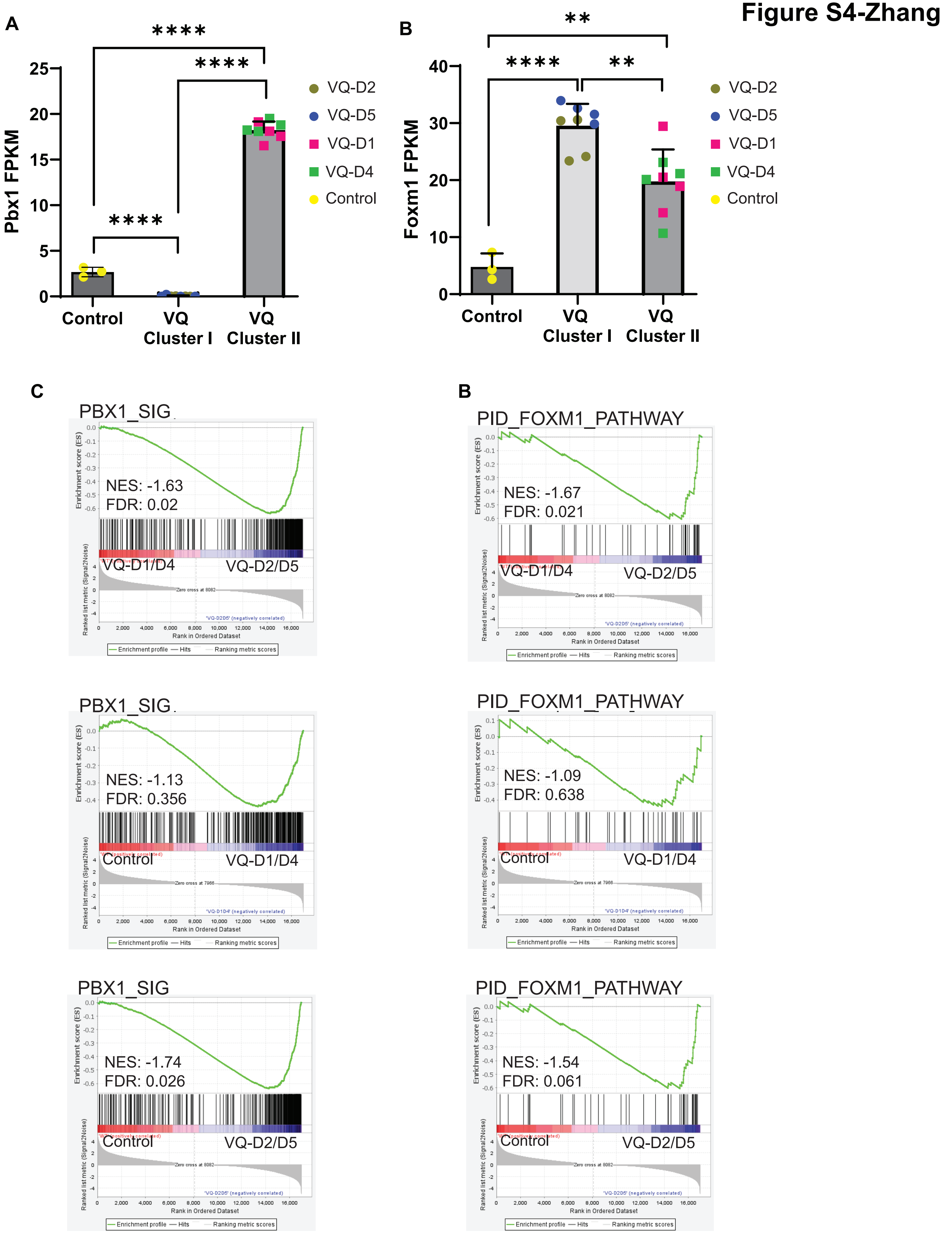
